# Development of an affinity-enhanced clinical candidate TCR targeting NY-ESO-1 with optimal potency and high specificity

**DOI:** 10.1101/2022.10.12.511904

**Authors:** Yusheng Ou, Min Liu, Kai Zhan, Wanli Wu, Jiao Huang, Hanli Sun, Hongjun Zheng, Xiaohong Yu, Haiping Gong, Zhaosheng Han, Aiyuan Chen, Youping Liao, Yanmei Lin, Ge Liu, Qiuping Liu, Ruijuan Ma, Yingyi Mei, Xianqing Tang, Zhiming Weng, Jieyi Wu, Yun Ye, Tingting Zhang, Jiehong Chen, Lin Chen, Qingfeng Ding, Hui Fan, Jiajia Hu, Jinhua Huang, Lihong Huang, Qiaowei Li, Siyun Li, Min Liu, Jia Shi, Xiangpeng Tan, Xiaoling Wang, Ruirui Xiang, Qingjia Yan, Jianbin Zhang, Wenjing Zheng, Shi Zhong

## Abstract

The clinical success of T-cell receptor (TCR)-based immunotherapy depends on the efficacy and specificity of TCRs. Naturally occurring TCRs have limited anti-tumor potency due to their low affinity for tumor antigens. Affinity enhancement is a promising strategy to generate highly potent TCRs. However, it is concerned that affinity-enhanced TCRs are prone to lose specificity. We isolated low affinity TCRs specific for NY-ESO-1_157-165_/HLA-A*02:01 from peripheral blood mononuclear cells of healthy donors. An affinity-enhanced TCR candidate with optimal affinity and specificity was generated using phage display and an extensive set of *in vitro* and *in vivo* assays. Alanine scanning mutagenesis showed that the TCR candidate retained specificity by making extensive contacts to the side chains of NY-ESO-1_157-165_ peptide. Adoptive transfer of T cells engineered with this candidate (termed TAEST16001) significantly inhibited tumor growth in subcutaneous, metastatic, and patient-derived xenograft (PDX) mouse tumor models. This study demonstrates that sophisticated engineering and screening techniques can be utilized to generate a clinical candidate TCR with potent anti-tumor activity without losing specificity. TAEST16001 was approved by the Center for Drug Evaluation (CDE) as the first TCR-based immunotherapy clinical trial in China (ClinicalTrials.gov Identifier: NCT03159585).

## 1 Introduction

By harnessing the immune system to fight cancer, immunotherapy has become the fourth pillar of cancer therapy, along with the conventional three pillars—surgery, radiation, and chemotherapy. As a prominent approach in immunotherapy, adoptive transfer of T cells genetically modified by T cell receptor (TCR) has demonstrated profound potential in clinical trials for both hematological malignancies and solid tumors.^1–4^ Nevertheless, the successful development of TCR-engineered T (TCR-T) cell therapy remains challenging owing to its efficacy and safety issues.

TCRs isolated from the peripheral T-cell repertoire typically have low affinity for tumor/self-antigens because highly self-reactive T cells are eliminated by the central tolerance mechanism. Those low affinity TCRs typically have low anti-tumor activities and are not suitable for clinical applications. Affinity enhancement through mutations in complementarity-determining regions (CDRs) can improve the functional activities of low avidity TCRs.^5–7^ Additionally, affinity-enhanced

TCRs can overcome certain immune escape mechanisms, one of which is the downregulation of antigen presentation,^8^ rendering low avidity T cells incapable of recognizing tumor cells. Affinity-enhanced TCRs show improved antigen sensitivity of T cells and hence may overcome this escape mechanism.^9^ Moreover, viral infections can escape immune responses by mutations in viral epitopes presented by human leukocyte antigen (HLA); high-affinity TCRs may circumvent this type of escape by effectively capturing those mutant epitopes.^10 11^

Safety concerns have been raised because of several adverse events or even fatal cases reported in TCR-T clinical trials.^12^ On-target/off-tumor toxicity arises when the cognate antigen of the transduced TCR is not restricted to the tumor. Gp100 and Mart-1 are differentiation antigens expressed in melanocytes. TCR-T cells targeting these antigens can effectively kill melanoma cells, but also lead to the destruction of melanocytes in healthy tissues, such as the eyes and skin.^13 14^ TCR-T cells may also cross-react with irrelevant self-antigens, typically containing epitopes similar in structure to the tumor epitope, and elicit unexpected off-target/off-tumor toxicity. This type of toxicity can be fatal to patients^15 16^ when irrelevant antigens are expressed in vital organs, such as Titin in heart tissues^17^ and Mage-A12 in the brain.^16^ Both on-target and off-target toxicities need to be carefully addressed during TCR-T development. Strategic *in vitro* screening techniques have also been proposed.^18 19^ Peptide libraries, such as alanine-scan, X-scan, and combinatorial peptide libraries, can be used to evaluate the propensity of the TCR to cross-react and even identify cross-reactive epitopes. In addition, functional assays of TCR-T cells against a large panel of healthy tissues are indispensable for identifying potential on- or off-target toxicity. Careful preclinical screening is required to mitigate unexpected toxicities before clinical trials.

The cancer testis (CT) antigen NY-ESO-1 has been recognized as an ideal immunotherapy target because it is highly immunogenic^21^ and is not expressed in normal tissues but is widely expressed in many tumor types, such as myxoid/round cell liposarcomas,^22^ synovial sarcoma,^23^ melanoma^24^ and, to a lesser extent, non-small cell lung cancer (NSCLC).^25^ There are various ongoing or finished NY-ESO-1 based cancer vaccine or adoptive cell transfer (ACT) clinical trials,^4^ among which adoptive transfer of NY-ESO-1 specific TCR-T cells showed the most promising results. The first clinical study conducted by the Rosenberg’s group achieved objective responses in four out of six synovial cell sarcoma patients and five out of eleven melanoma patients^26^ using the affinity-enhanced TCR 1G4-α95:LY. A follow-up trial with additional patients was conducted, and the overall response rates combining both cohorts were 61% for patients with synovial cell sarcoma and 55% for patients with melanoma.^3^ Another phase I/II trial conducted by June et al. using the NY-ESO^c259^ TCR (which is identical to 1G4-α95:LY in sequence) reported an 80% response rate and 19.1-month progression-free survival for patients with multiple myeloma.^2^ These clinical trials are of great significance. First, they demonstrated substantial clinical benefits for solid tumors in non-melanoma patients. Second, the NY-ESO-1 specific TCRs used in both groups were affinity enhanced from the original wild 1G4 TCR,^21 27^ rather than the wild-type TCRs used in previous ACT trials. Finally, no toxicity was observed, even for enhanced affinities. In contrast, in melanoma trials, higher affinity TCRs targeting melanoma antigens Mart-1 or gp100 led to on-target/off-tumor toxicity,^13^ which was not observed in a related trial with a lower affinity TCR,^28^ suggesting that NY-ESO-1 has a better safety profile than melanoma antigens.

Although affinity-enhancement of TCRs can significantly improve their anti-tumor avidity, substantial challenges regarding specificity must be addressed. Here, we describe the development of TAEST16001, a clinically proven affinity-enhanced NY-ESO-1-specific TCR-T therapy. We demonstrate that affinity-enhancement does not alter TCR specificity. Together with other IND-enabling studies, these results led to the first approved TCR-T clinical trial by the Center for Drug Evaluation (CDE) in China (ClinicalTrials.gov Identifier: NCT03159585).

## 2 Methods

### 2.1 Cells

Peripheral blood mononuclear cells (PBMCs) were prepared from buffy coats of healthy blood donors (Guangzhou Blood Center, Guangzhou, China) using Lymphoprep™ (STEMCELL Technologies) density gradient centrifugation according to the manufacturer’s instructions. CD8^+^ and CD3^+^ T cells were isolated from PBMCs using the EasySep™ Human CD8+ T Cell Isolation Kit and EasySep™ Human T Cell Isolation Kit (both from STEMCELL Technologies), respectively. T cells specific for NY-ESO-1_157-165_ were isolated from HLA-A*02:01^+^ PBMCs using tetramer-guided sorting on a BD AriaIII cell sorter, cloned, and expanded as previously described.^29^ T2, A375, IM9, MDA-MB-231, NCI-H1299, and K562 cells were obtained from the American Type Culture Collection (ATCC), and cultured in RPMI 1640 (Gibco, T2, IM9, NCI-H1299), DMEM (Gibco, A375), L15 (Gibco, MDA-MB-231) or IMDM (Gibco, K562) medium supplemented with 10% heat-inactivated fetal bovine serum (Gibco). NY-ESO-1 or HLA overexpressing cell lines were established using lentiviral transduction. The lung cancer cell line A549 expressing luciferase was purchased from Shanghai Model Organisms Center, Inc. and overexpressed with HLA-A*02:01 and NY-ESO-1, generating A549^Luciferase/A0201/NY-ESO-1^. The NSCLC PDX model LU0367 was established by Crown Bioscience Inc. and confirmed as HLA-A*02:01^+^ and NY-ESO-1^+^ using RNAseq and immunohistochemistry staining. Lymphoblastoid cell lines (LCLs) were established from the PBMCs of different healthy donors via Epstein-Barr virus (EBV) transformation. All normal primary cells were obtained from ScienCell Research Laboratories and were cultured according to the instructions.

### 2.2 Peptides, peptide-major histocompatibility complex (pMHC), and tetramer production

All peptides were synthesized at >95% purity by GenScript (Jiangsu, China), and verified using mass spectroscopy. As previously described, biotinylated pMHCs and tetramers were produced in-house.^11^

### 2.3 TCR gene cloning

Total RNA was extracted from the T cells using the RNeasy Mini RNA Isolation Kit (Qiagen). TCR α- and β-chain genes were reverse transcribed and amplified from the RNA using the SMARTer RACE Kit (Clontech Laboratories, Inc). The amplified TCR genes were sub-cloned into the pEF- 1α/pENTR vector (Addgene) and sequenced.

### 2.4 mRNA preparation and electroporation

The constant regions of the human TCR genes were replaced with murine constant genes by overlapping PCRs as previously described,^14^ and the human/murine hybrid TCR genes were sub-cloned into the pGEM-4Z vector (Promega Corporation) for mRNA expression. mRNAs encoding TCR genes were transcribed *in vitro* using the mMESSAGE mMACHINE® T7 Ultra Kit (Life) using the Avr□ (NEB) linearized plasmid DNA as templates. The transcribed mRNAs were purified with the RNeasy Mini Kit (Qiagen) and stored in −80 °C freezer.

For activation, T cells were mixed with Dynabeads Human T-Activator CD3/CD28 (Gibco) at a 2:1 ratio in a 24 well plate in RPMI 1640 medium (Gibco) supplemented with 10% heat-inactivated fetal bovine serum (Gibco) and 100 IU/mL IL-2 (Beijing Four Rings Biopharmaceuticals Co., Ltd.) and cultured for 3 days in a 37 °C/5% CO_2_ incubator. *In vitro* transcribed mRNAs encoding the TCR α- and β-chains were mixed at a 1:1 ratio and electroporated in activated T cells on a Lonza 4D-Nucleofector^TM^ device using the P3 Primary Cell 4D-Nucleofector™ X Kit (Lonza) according to the recommendations of the manufacturer. After electroporation, the cells were transferred to a fresh medium and cultured in a 37 °C incubator.

### 2.5 TCR affinity engineering and affinity measurement

Phage display screening was employed to engineer TCR molecules, as previously described.^27^ Briefly, phage display libraries were constructed in the CDR3 regions of the α- and β-chains of the TCR by random mutagenesis at a span of four–five amino acids. High-affinity TCR variants were selected by panning CDR3 phage libraries on immobilized pMHC (NY-ESO-1 _157-165_/HLA-A*02:01). After three rounds of panning, phage clones were picked and tested for their binding to pMHC by inhibitive phage ELISA,^27^ and positive clones were sequenced. Disulfide bond-linked soluble TCRs were produced as previously described.^11^ The binding between the soluble TCR and pMHC was determined using surface plasmon resonance (SPR) on a Biacore T200 (GE Healthcare) as previously described.^11^

### 2.6 Lentiviral transduction

The α- and β-chains of codon-optimized TCR genes were linked with a P2A self-cleavage peptide sequence, synthesized (Genscript) and sub-cloned into a self-inactivating lentiviral vector under the EF1-a promoter. Lentiviral particles encoding TCRs were produced in-house using the 3^rd^ generation lentivirus packaging system. Peripheral blood leukocytes (PBLs) were transduced with lentiviral particles at a multiplicity of infection (MOI) of 5 for *in vitro* assays and an MOI of 1 for *in vivo* experiments.

### 2.7 Flow cytometry analysis

Anti-CD3 (clone OKT3), anti-CD8 (clone RPA-T8), anti-TCR Vβ8 (clone JR2), and anti-mTRBC (clone H57-597) antibodies were purchased from Biolegend (San Diego, CA). Cells were analyzed on a Guava easyCyte 12HT cytometer (Millipore).

### 2.8 Cellular assays

Enzyme-linked immunospot (ELISpot) assays were conducted using the Human IFN-γ ELISpot Set (BD Biosciences) according to the manufacturer’s instructions. A mixture of 2000 effector cells (TCR-T cells), 20000 target cells (T2 or other cell lines), and peptides (only for T2 cells) were co-cultured overnight at 37 °C and 5% CO_2_ in an ELISpot plate coated with a capture antibody. The spots were counted using an AID ELISPOT READER SYSTEM (Autoimmun Diagnostika GmbH).

Cytotoxicity was determined by the lactate dehydrogenase (LDH) release-based assay using the CytoTox 96® non-radioactive cytotoxicity assay kit (Promega Corporation) as previously described.^11^ Live-cell imaging assays were performed using the IncuCyte platform (Essen BioScience). Briefly, tumor cells were plated at 10,000 cells per well in a 96-well plate (Corning) and incubated overnight at 37 °C and 5% CO_2_ in R10 without phenol red (Gibco). The following day, cells were washed twice and cultured in R10 (without phenol red) in the presence of T cells at a 1:1 effector:target cell ratio in the presence of the caspase-3/7 Green Detection Reagent (Invitrogen). Images were taken every 2 h at 10× magnification. The number of apoptotic cells per mm^2^ was quantified using the IncuCyte ZOOM software.

### 2.9 Xenograft models

NOD/SCID/IL2gR^-/-^ (NSG) mice aged 6–8 weeks were purchased from Biocytogen (Beijing, China) and maintained under sterile environmental conditions in a 12 h light/dark cycle. The Institutional Animal Care Committee approved all experimental procedures.

For the subcutaneous tumor model, NSG mice were injected subcutaneously into a single flank with 1 × 10^7^ NCI-H1299-A2 cell line or 3 × 3 × 3 mm^3^ patient-derived xenograft (PDX) fragments. Tumor sizes were measured twice per week with calipers in two perpendicular dimensions, and tumor volumes were calculated using the following formula: volume (mm^3^) = (length × width^2^)/2. When the tumor volume reached a size of 80–100 mm^3^, TAEST16001 or control cells were administered via tail vein injection. At the same time, 50000 IU IL-2 was administered intraperitoneally every 24 h for 5 days.

For the lung metastatic model, 3 × 10^6^ A549^Luciferase/A0201/NY-ESO-1^ were injected intravenously into NSG mice. Mice were monitored weekly for tumor growth by bioluminescence imaging of anesthetized mice using a Bruker In-Vivo Xtreme system. For imaging, 10 mg/kg D-luciferin re-suspended in sterile PBS at a 15 mg/mL concentration was administered intraperitoneally. Mice were imaged 5 min after luciferin injection, and serial images were collected under X-ray or fluorescent light. Data were analyzed using the Bruker molecular imaging software using images taken with identical settings for mice in each group at each time point. Imaging data were converted to net photons/mm^2^ for the quantitative analysis.

## 3 RESULTS

### 3.1 Isolation and characterization of NY-ESO-1_157-165_/HLA-A*02:01 specific TCRs from PBMCs of healthy donors

NY-ESO-1 _157-165_/HLA-A*02:01-specific T cell clones were isolated from PBMCs of HLA-A*02:01 (HLA-A2) positive healthy donors. TCR genes were amplified from the clones using 5’ rapid amplification of cDNA ends (5’ RACE). Three unique TCRs (SL1, SL2 and SL3, table 1) were selected for the subsequent affinity engineering. Soluble TCRs were generated through *in vitro* refolding, and their binding kinetics to soluble NY-ESO-1_157-165_/HLA-A2 complex was determined using surface plasmon resonance (SPR). The binding affinities of the TCRs (43–250 μM, table 1) are in the physiological affinity range of wild-type TCRs specific for tumor-associated self-antigens.^30^ To verify the function of the three TCRs, mRNA encoding the murinized TCR genes (i.e. the substitution of the murine constant domains for the human ones)^27^ were electroporated into CD8^+^ T cells activated by anti-CD3/CD28 beads. The surface expression levels of the three TCRs were comparable, as evidenced by similar staining (>90%) using anti-murine TCR β-chain antibody (figure 1A-C right panels). However, the tetramer binding levels were highly variable: SL1, SL2 or SL3-transduced T cells showed weak, intermedia or high tetramer staining (figure 1A-C left panels), respectively, consistent with the relative affinities of the three TCRs (SL1 *K*_D_ = 250 μM, SL2 *K*_D_ = 55 μM and SL3 *K*_D_ = 43 μM, table 1). The functional activities of the three TCRs were also consistent with their binding affinities. T cells expressing SL3 (the highest affinity of the three TCR) showed the most potent functional activity to T2 cells pulsed with NY-ESO-1 _157-165_ peptide in a concentration-dependent manner, as determined using the IFN-γ ELISpot assay (figure 1F). The functional activity was much weaker for the lower affinity TCR SL2 (*K*_D_ = 55 μM, figure 1E) and almost undetectable for SL1 (*K*_D_ = 250 μM, figure 1D).

**Table 1.**
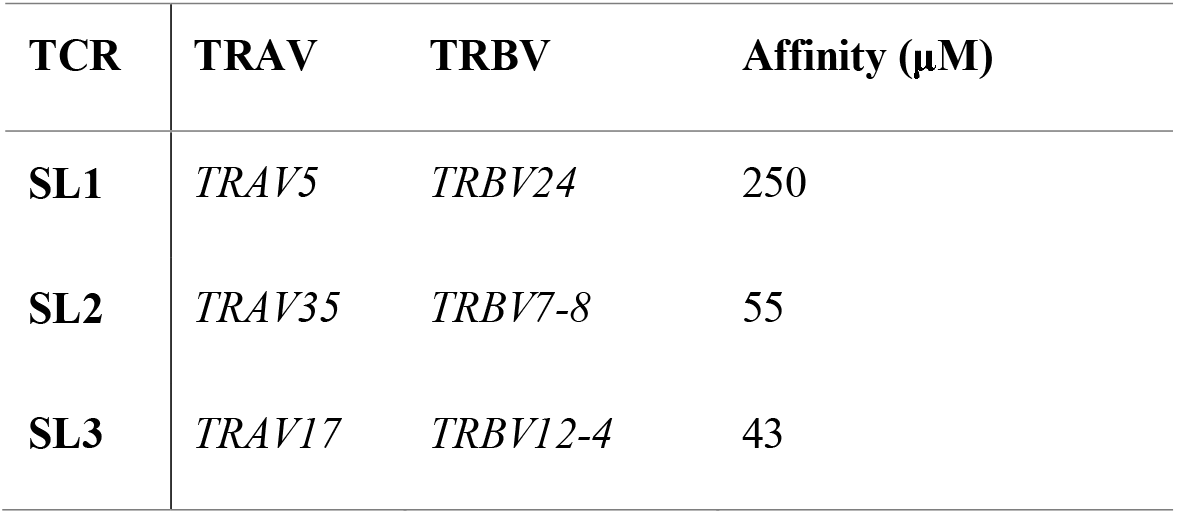
Three NY-ESO-1 _157-165_/HLA-A2-specific TCRs isolated from health donors. The binding affinity of each TCR to its cognate ligand was determined by SPR analysis.

**Figure 1.**
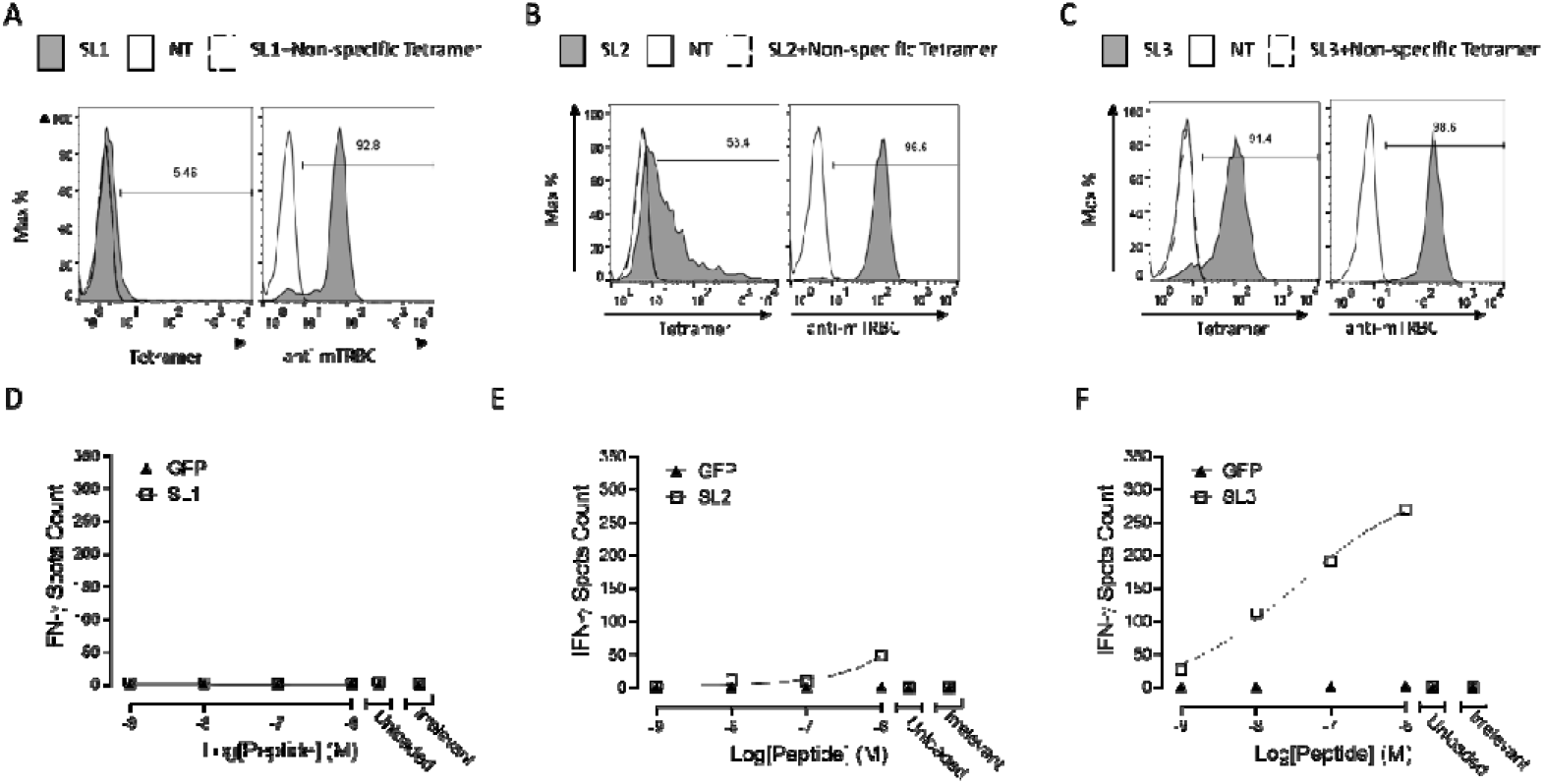
Characterization of three TCRs specific for NY-ESO-1 157-165/HLA-A2. mRNAs encoding murinized SL1 (A), SL2 (B) or SL3 (C) TCR genes were electroplated in CD8^+^ T cells activated by anti-CD3/CD28 beads, and the expression of each TCR was evaluated by flow cytometry using anti-murine TCR-β antibody, NY-ESO-1 _157-165_/HLA-A2 tetramer or non-specific tetramer staining. Non-transduced T cells (NT) were used as a negative control. (B) CD8^+^ T cells electroplated with mRNA encoding SL1 (D), SL2 (E) or SL3 (F) TCR genes co-cultured with T2 cells loaded with NY-ESO-1 157-165, an irrelevant peptide (10^-6^ M gp100280-288, Irrelevant) or no peptide (Unloaded), and IFNγ release was determined using the IFN-γ ELISpot assay. GFP transduced T cells served as a negative control. Data indicate mean+/-SD of triplicates.

### 3.2 Affinity enhancement by phage display

We selected SL2 and SL3 for affinity enhancement and excluded SL1 from further development because of its low affinity (*K*_D_ = 250 μM) and poor functional activity (figure 1D). TCRs were displayed on the phage surface by fusing them to the gene III product of M13 phage,^27^ and mutant libraries were generated by introducing mutations in the complementarity-determining region 3 (CDR3) regions of both the α- and β-chains. Several rounds of phage screening using immobilized NY-ESO-1 _157-165_/HLA-A2 yielded many unique mutants. In total, 12 SL2 mutants (5 α-chain and 7 β-chain) and 13 SL3 mutants (6 α-chain and 7 β-chain) were produced as soluble TCRs, and their binding kinetics to NY-ESO-1_157-165_/HLA-A2 were measured using SPR. We obtained mutants with a wide range of affinity enhancement for both TCRs (table 2). The affinities of the SL2 mutants ranged from ~8.8 μM to ~0.5 μM (6-fold to 105-fold increase compared to the wild-type TCR, *K_D_* = 55 μM). For SL3, the affinity range of the mutants is between ~1.7 μM and ~0.1 μM (25 to 383-fold increase compared to the wild-type TCR, *K_D_* = 43 μM).

**Table 2.**
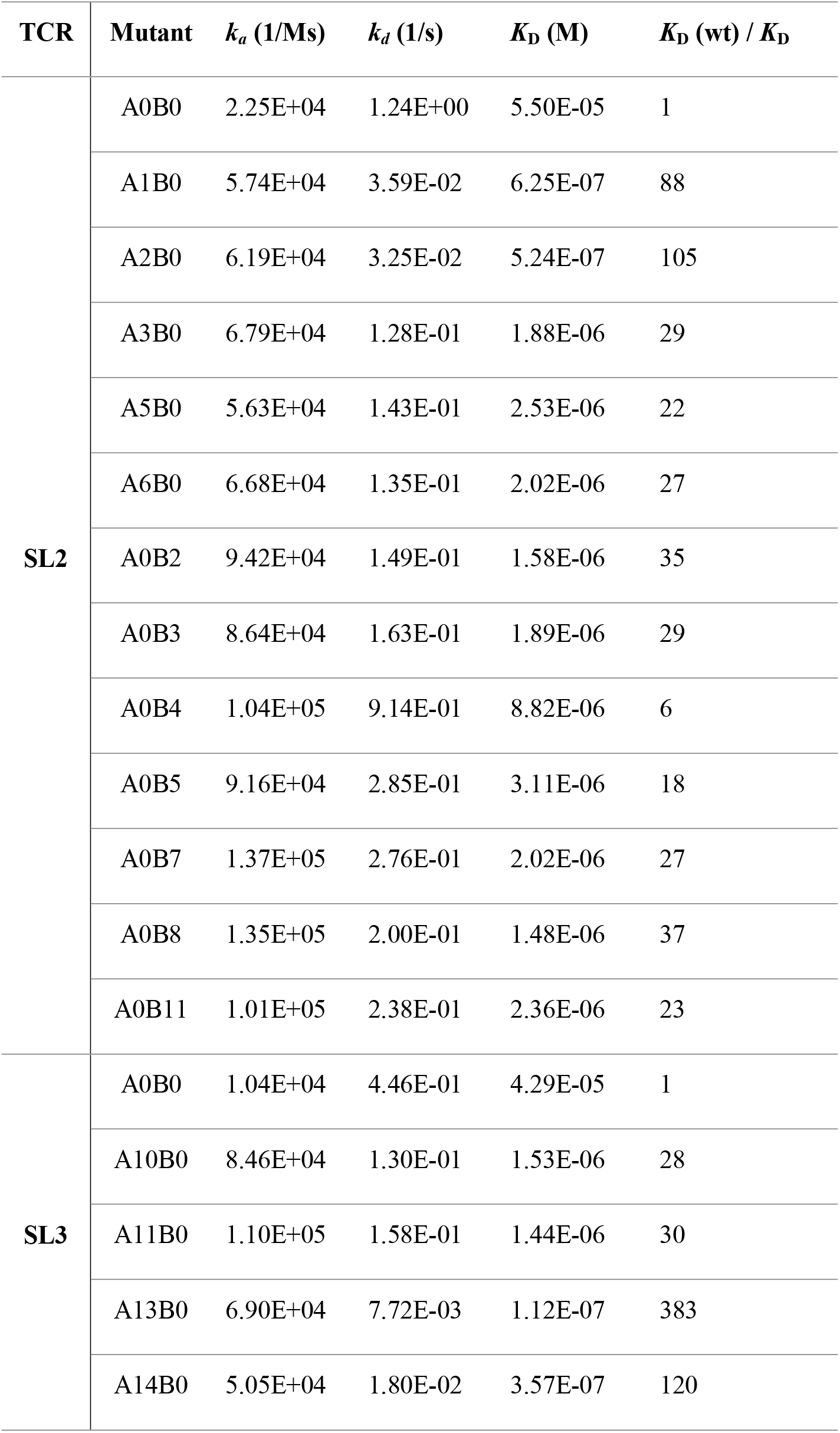
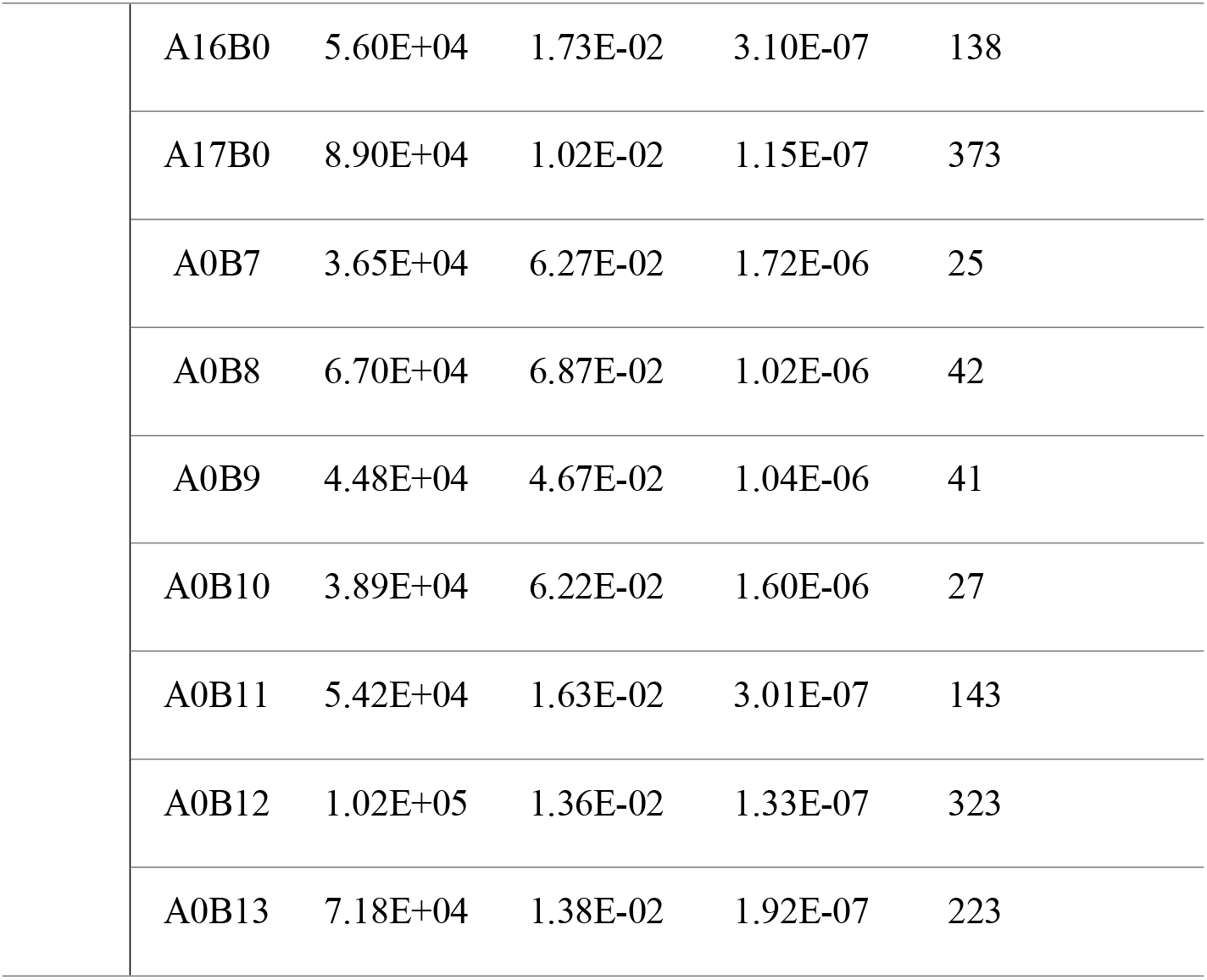
Binding properties of SL2 and SL3 TCR affinity-enhanced mutants generated using phage display. A0B0, AXB0 and A0BX represent wild-type (wt), *α* -chain mutant and β-chain mutant, respectively. The affinity and kinetic rate constants of each mutant were determined by SPR. The fold increase in affinity over the wt TCR was calculated using equation: *K*_D_ (wt)/*K*_D_.

### 3.3 Functional screening of affinity-enhanced TCR mutants

To screen the functional avidity of the SL2 and SL3 mutants, peripheral blood leukocytes (PBLs) electroporated with mRNA encoding the murinized TCR genes were co-cultured with T2 cells pulsed with a titration of NY-ESO-1_157-165_. 1G4-α95:LY was used as a reference TCR and green fluorescent protein (GFP) as a negative control in the screening experiments. The functional activities of the mutants were evaluated using IFN-γ ELISpot assays and compared with the reference TCR. Our goal was to identify TCR mutants demonstrating equal or higher functional potency than the reference

TCR without losing specificity. The SL2 α-chain mutants showed enhanced functional potency compared to the wild-type SL2 (SL2-A0B0), and still retained specificity, but none were as potent as the reference TCR (online supplemental figure 1A). The mutations on the SL2 β-chain led to inferior specificity, as shown by excessive release of IFN-γ after pulsing with a non-specific peptide or no peptide (online supplemental figure 1B). Therefore, none of the SL2 mutants we screened satisfied our goal. We continued to assess the SL3 mutants and identified two mutants (SL3-A10B0 and SL3-A0B9) with potency comparable to that of the reference TCR, and without apparent non-specific activations (online supplemental figure 2A and 2B). Other SL3 mutants showed either lower potency (such as SL3-A14B0) or non-specificity (such as SL3-A16B0). However, on close examination of SL3-A0B9, we found a slightly higher than background activation when no peptide was pulsed (online supplemental figure 2B SL3-A0B9 unloaded), suggesting potential non-specificity. To investigate this observation further, we tested the activation SL3-A0B9 or SL3-A10B0-transduced T cells against tumor cell lines using IFN-γ or Granzyme B ELISpot assays, and found higher than background activation for SL3-A0B9, but not for SL3-A10B0 (online supplemental figure 2C). Therefore, SL3-A0B9 was excluded from further development due to its non-specificity.

We further evaluated SL3-A10B0-transduced T cells against a panel of antigen-expressing tumor cell lines that naturally process and present antigenic peptides. The expression levels of NY-ESO-1 and NY-ESO-2 (which also contains the SLLMWITQC epitope) in tumor cell lines were assessed using the NanoString nCounter system (online supplemental table 1). We identified tumor cell lines expressing either NY-ESO-1 (A375, NCI-H1299), or NY-ESO-2 (K562, NCI-H522), or both (IM9, U266B1). IFN-γ (figure 2A) or Granzyme B (figure 2B) secretion was detected upon co-culture of SL3-A10B0-transduced T cells with HLA-A2 and NY-ESO-1/2 double-positive tumor cell lines (either wild-type or HLA-A2/antigen overexpressed), but not with HLA-A2 or NY-ESO-1/2 negative cell lines (figure 2A). The amount of IFN-γ or Granzyme B secretion by SL3-A10B0-transduced T cells was significantly higher than that by SL3-A0B0 (wild-type SL3)-transduced T cells, and comparable to that by 1G4-α95:LY-transduced T cells, suggesting that SL3-A10B0 is an affinity-enhanced TCR with superior functional avidity.

**Figure 2.**
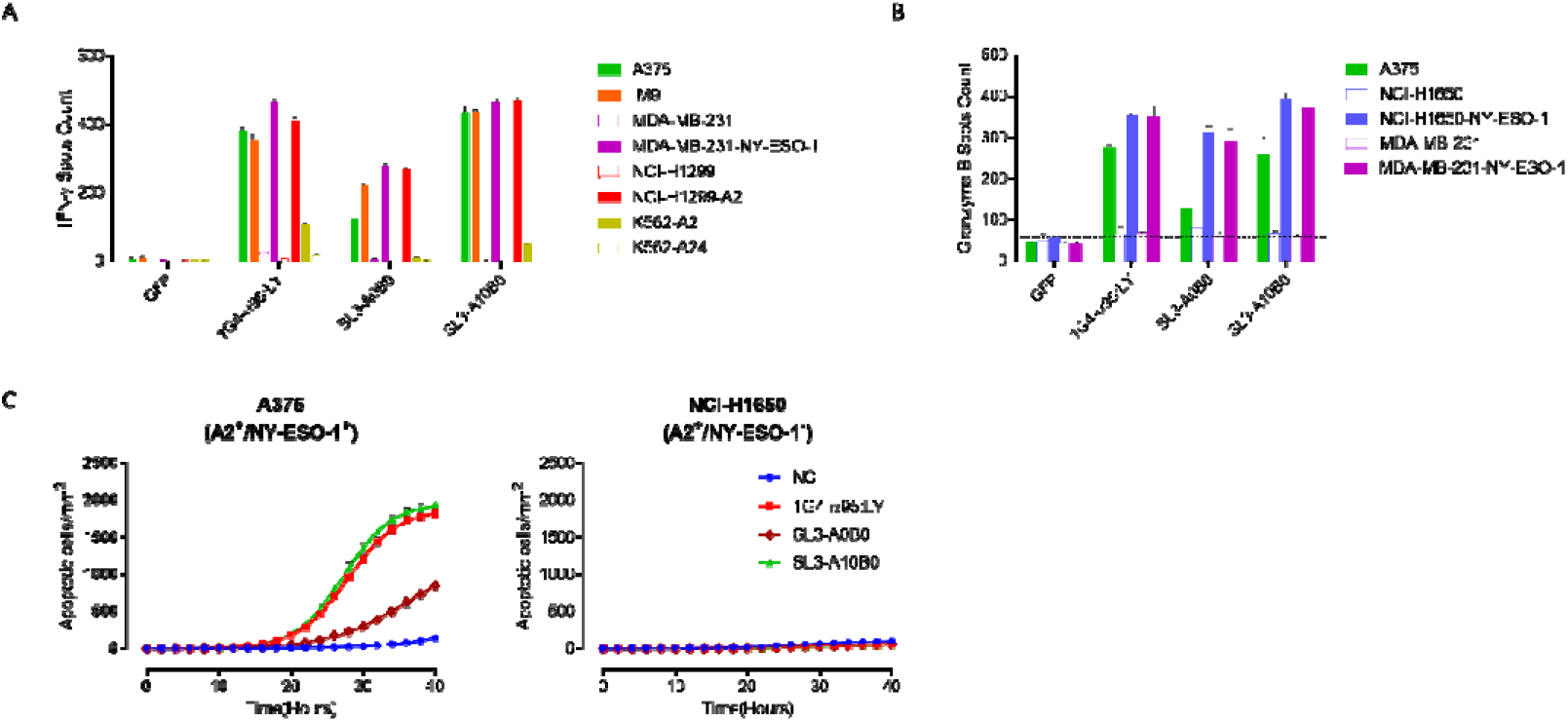
Affinity-enhanced SL3 TCR mutant (SL3-A10B0) shows superior functional avidity. IFN-γ release (A) or Granzyme B (B) release of CD8^+^ T cells expressing SL3-A0B0 (SL3 wild-type TCR) or SL3-A10B0 after co-culturing with tumor cell lines. T cells expressing GFP served as a negative control and 1G4-α95:LY as a positive control. The expression of NY-ESO-1 and NY-ESO-2 of the cell lines was determined using the NanoString nCounter Analysis (online supplemental table 1): A375 (HLA-A2^+^, NY-ESO-1^+^/NY-ESO-2^-^), IM9 (HLA-A2^+^, NY-ESO-1^+^/NY-ESO-2^+^), MDA-MB-231 (HLA-A2^+^, NY-ESO-17NY-ESO-2^-^), MDA-MB-231-NY-ESO-1 (HLA-A2^+^, NY-ESO-1 overexpressing), NCI-H1299 (HLA-A2^-^, NY-ESO-1^+^/NY-ESO-2^-^), NCI-H1299-A2 (HLA-A2 overexpressing, NY-ESO-1^+^/NY-ESO-2^-^), K562-A2 (HLA-A2 overexpressing, NY-ESO-1^-^/NY-ESO-2^+^), K562-A24 (HLA-A2^-^, NY-ESO-1^-^/NY-ESO-2^+^), NCI-1650 (HLA-A2^+^, NY-ESO-1^-^/NY-ESO-2^-^), NCI-1650-NY-ESO-1 (HLA-A2^+^, NY-ESO-1 overexpressing). (C) The lysis of tumor cells mediated by T cells expressing SL3-A0B0 or SL3-A10B0 using kinetic live cell imaging assay. None-transduced T cells (NC), SL3-A10B0 or SL3-A0B0 expressing T cells were co-cultured with tumor cells A375 (HLA-A2^+^, NY-ESO-1^+^, left panel) or NCI-H1650 (HLA-A2^+^, NY-ESO-1^-^, right panel) at 1:1 ratio in the presence of the caspase-3/7 green detection reagent and images (10× magnification) were captured every 2 h for 40 h in an IncuCyte® ZOOM system. Representative images are shown in online supplemental figure 3. The number of apoptotic tumor cells was measured in the IncuCyte® ZOOM software using green object counting. Data indicate mean+/-SD of triplicates.

Next, we studied the dynamic killing of tumor cell lines using the Incucyte Live Cell Imaging System, enabling visualization of caspase 3/7-dependent apoptosis in real-time (figure 2C and online supplemental figure 3). The killing of A375 cells (HLA-A2^+^, NY-ESO-1^+^) was observed at approximately 15 h. SL3-A10B0- and 1G4-α95:LY-transduced T cells showed a significantly higher rate of killing than SL3-A0B0-transduced. No non-specific killing of antigen-negative cells (NCI-H1650, HLA-A2^+^/NY-ESO-1^-^) was observed in the course of the measurements (figure 2C and online supplemental figure 3 right panel). Collectively, SL3-A10B0 was verified as a high avidity TCR and was selected as our lead candidate for further investigation. The affinity of SL3-A10B0 was determined to be ~1.5 μM (online supplemental figure 4), a ~28-fold increase compared to that of the wild-type (~43 μM).

### 3.4 Assessment of *in vitro* efficacy of TAEST16001

Codon-optimized SL3-A10B0 gene was cloned into a lentiviral vector. SL3-A10B0-tranduced T cells were produced using the 3^rd^ generation lentivirus-based gene transfer system. T cells transduced with the SL3-A10B0 lentiviral vector were designated TAEST16001, where TAEST stands for TCR affinity-enhanced specific T cells. TAEST16001 showed high levels of TCR expression (>85%), as determined by tetramer and anti-human Vβ8 (specific for the variable region of SL3-A10B0 β-chain) staining (figure 3A). TAEST16001 mediated specific IFN-γ release when co-cultured with HLA-A2 and NY-ESO-1/2 double positive cell lines, but not with HLA-A2 or NY-ESO-1/2 negative cell lines (figure 3B). TAEST16001 also induced specific killing of A375 (HLA-A2^+^ and NY-ESO-1^+^), but not NCI-H1650 (HLA-A2^+^ and NY-ESO-1^-^) (figure 3C). Taken together, these data indicate superior *in vitro* anti-tumor potency of TAEST16001.

**Figure 3.**
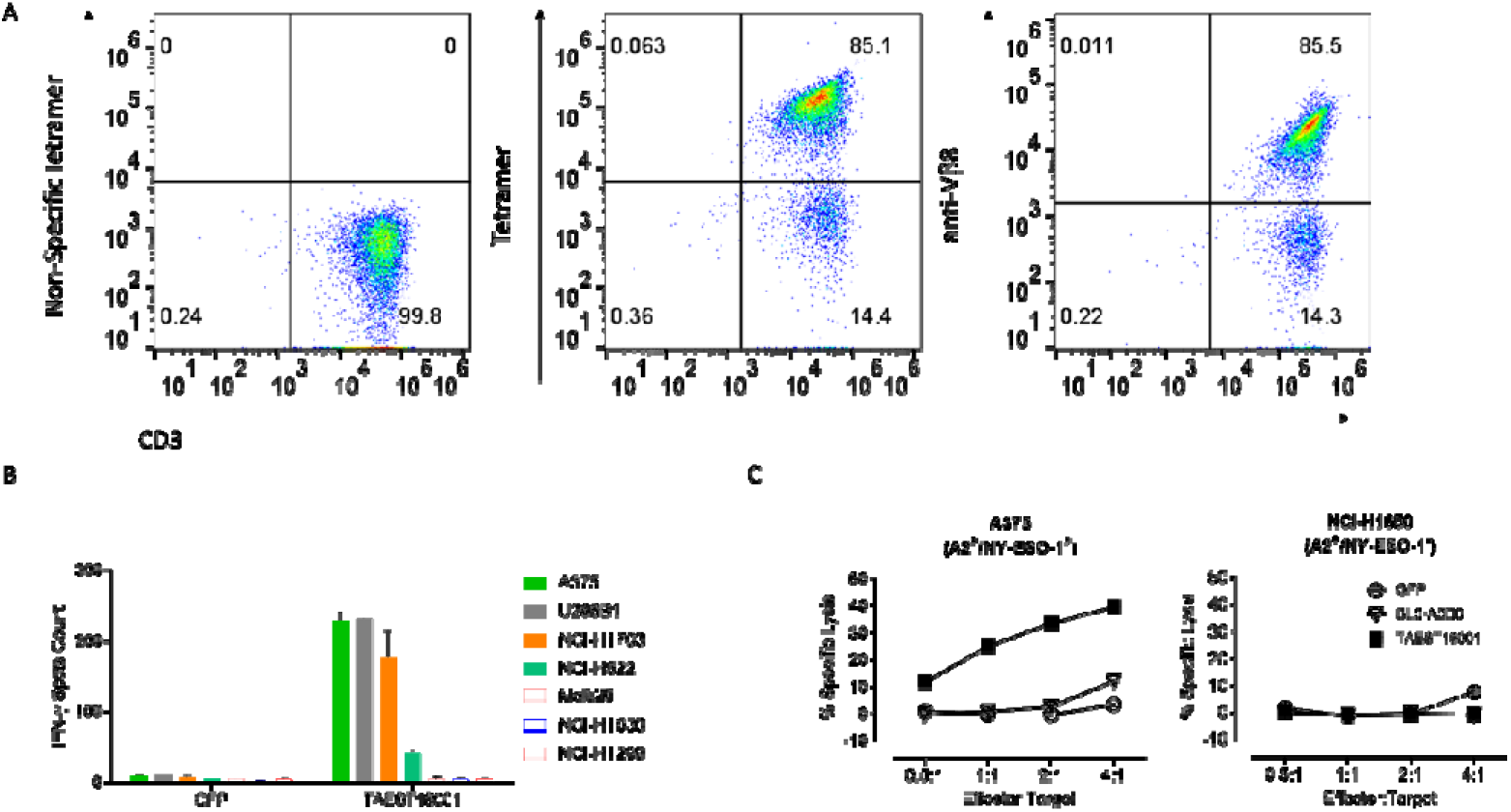
TAEST16001 shows high anti-tumor efficacy *in vitro.* (A) The TCR surface expression of TAEST16001 was assessed using flow cytometry. The cells were double stained for anti-CD3 antibody together with NY-ESO-1157-165/HLA-A2 tetramer, anti-Vβ antibody, or an irrelevant non-specific tetramer as a control. (B) IFN-γ release of TAEST16001 after co-culturing with tumor cell lines. T cells expressing GFP served as a negative control. The expression of NY-ESO-1 and NY-ESO-2 of the cell lines was determined using the NanoString nCounter Analysis (online supplemental table 1): A375 (HLA-A2^+^, NY-ESO-1^+^/NY-ESO-2^-^), U266B1 (HLA-A2^+^, NY-ESO-1^+^/NY-ESO-2^+^), NCI-H1703 (HLA-A2^+^, NY-ESO-1^+^/NY-ESO-2^-^), NCI-H522 (HLA-A2^+^, NY-ESO-1^-^/NY-ESO-2^+^), MEL526 (HLA-A2^+^, NY-ESO-1TNY-ESO-2^-^), NCI-H1650 (HLA-A2^+^, NY-ESO-1TNY-ESO-2^-^) and NCI-H1299 (HLA-A2^-^, NY-ESO-1^+^/NY-ESO-2^-^). (C) TAEST16001, SL3-A0B0 (for comparison), or GFP (as a negative control) transduced T cells were co-cultured with A375 (left panel) or NCI-H1650 (right panel) at the indicated effector:target ratios for 24 h and the specific killing of tumor cells was assessed using the LDH release assay. Data indicate mean+/-SD of triplicates.

### 3.5 Assessment of the *in vitro* safety profile of TAEST16001

To mitigate this risk of potential cross-reactivity of TAEST16001, we applied several *in vitro* strategies. First, the binding of SL3-A10B0 soluble protein to a panel of irrelevant peptide-HLA-A2 complexes (online supplemental table 2) was determined using SPR measurements. No detectable non-specific binding was observed for any of the complexes, suggesting that affinity enhancement did not change binding specificity.

Next, we investigated the binding patterns of SL3-A10B0 and 1G4-α95:LY using the alanine-scanning mutagenesis strategy (each of the amino acids of NY-ESO-1 _157-165_, except for the anchor residue at position 2, was sequentially replaced by alanine)^17 32^. To determine the effect of alanine substitutions on functional activities of TCRs, IFN-γ release of SL3-A10B0- or 1G4-α95:LY-transduced T cells upon co-culture with T2 cells pulsed with wild-type and mutant peptides was assessed using ELISpot assays (figure 4A). To determine the effect of mutation on binding kinetics, the binding of soluble SL3-A10B0 and 1G4-α95:LY to alanine-substituted peptide-HLA complexes was analyzed using SPR (online supplemental table 3). The changes in binding affinities to mutant peptide-HLA relative to wild-type peptide-HLA were calculated (figure 4A bottom table). For 1G4-α95:LY, alanine mutations at positions 1, 3, 7 and 9 had a minor effect on both binding affinities (<14-fold) and functional activities; mutations at positions 4, 6 and 8 led to a modest decrease in affinities (~50 to ~120-fold) and functional activities; mutation at position 5 abrogated TCR binding and functional activity. These results are in agreement with the crystal structure of wild-type 1G4 TCR in complex with NY-ESO-1 _157_._165_/HLA-A2^33^ and alanine-scanning mutagenesis studies of wild-type 1G4 TCR.^34^ For SL3-A10B0, mutations at 1, 8 and 9 had a minor effect on both binding affinities (<1.2-fold) and functional activities; mutation at position 4 led to a modest decrease in affinity (~120-fold) and functional activities; mutations at positions 3, 5, 6 and 7 significantly reduced affinities (>289-fold) and abrogated functional activities. In summary, we found that one residue (position 5) was critical and three were less critical (positions 4, 6 and 8) for 1G4-α95:LY binding to NY-ESO-1_157-165_/HLA-A2, whereas four residues (positions 3, 5, 6 and 7) were critical and one (position 4) was less critical for SL3-A10B0 binding to NY-ESO-1_157-165_/HLA-A2. Our results indicate that SL3-A10B0 has a higher level of specificity than 1G4-α95:LY and thus is less likely to cross-react.

**Figure 4.**
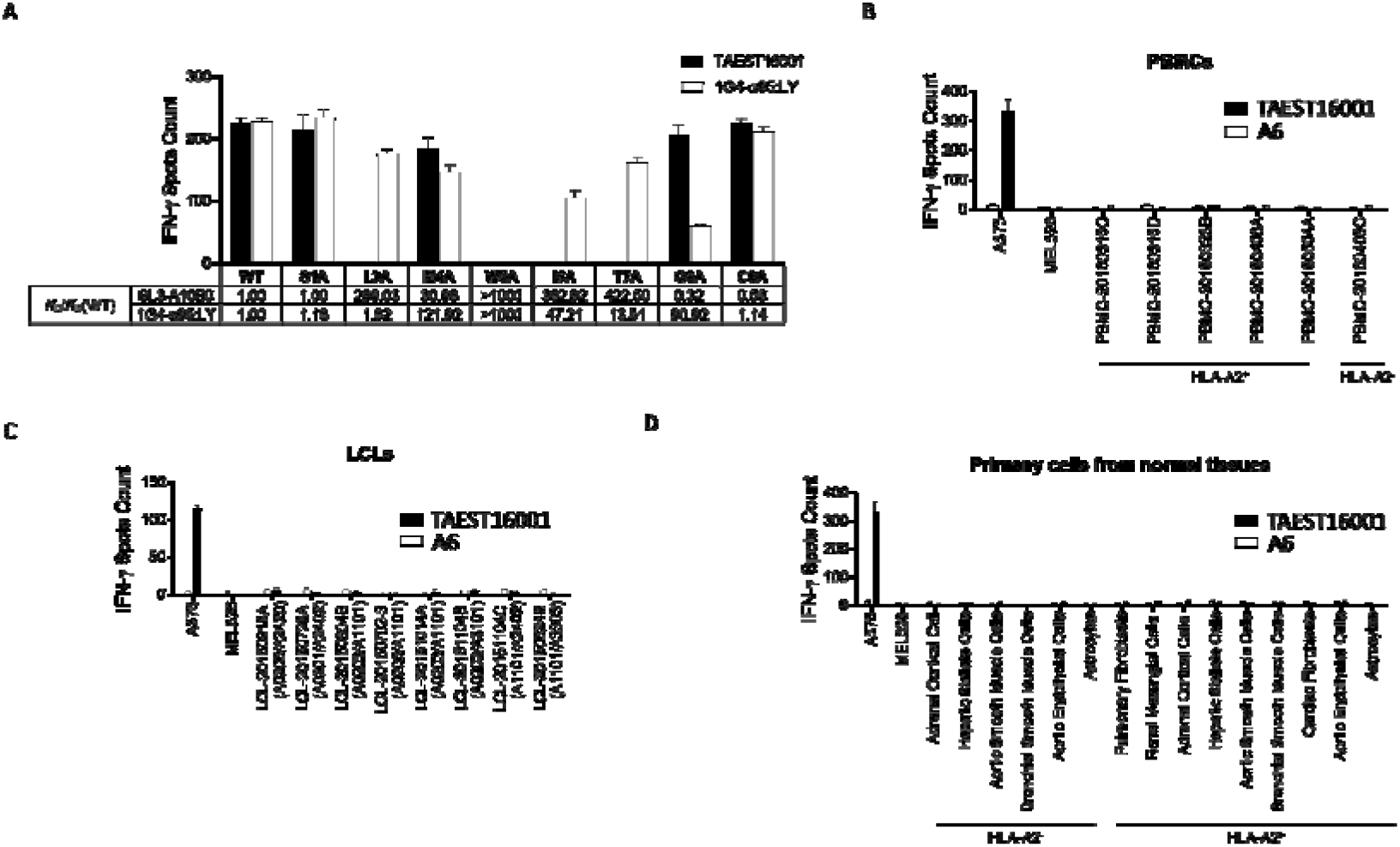
Specificity analysis of TAEST16001. (A) Alanine-scanning analysis. Each residue of NY-ESO-1 157-165 was substituted with alanine, except for the anchor residue at position 2. Upper panel: T2 cells loaded with alanine-substituted peptides co-cultured with TAEST16001 or 1G4-α95:LY-transduced T cells. IFN-γ release was assessed using the ELISpot assay. Lower panel: The binding properties of SL3-A10B0 or 1G4-α95:LY to alanine-substituted peptide-HLA-A2 complexes were analyzed using SPR (online supplemental table 3). The *K*_D_ values of the TCRs binding to mutant peptide-HLAs, relative to binding to the WT peptide-HLA [*K*_D_/*K*_D_(WT)] were calculated. (B-D) Non-specific activation of TAEST16001 by a panel of PBMCs isolated from healthy donors (B), lymphoblastoid cell lines (LCLs, C, HLA typing was indicated), and primary cells derived from normal tissues (D). A375 and MEL526 tumor cell lines were used as positive and negative target cell controls, respectively. A6 (mock TCR)-transduced T cells were included as effector cell control. The non-specific activation was determined using the ELISpot assay. Data indicate mean+/-SD of triplicates.

To further investigate whether TAEST16001 has potential off-target/off-tumor reactivity, we performed extensive *in vitro* analysis on several panels of normal cells: PBMCs from six donors (five of which were HLA-A2^+^, Fig 4B), LCLs derived from eight donors (Fig 4C), and a set of fifteen normal tissue-derived primary cells (nine of which were HLA-A2^+^, Fig 4D). Using IFN-γ ELISpot assays as a readout for T cell activation, no activity was observed against any of these cells, suggesting that off-target toxicity is not a concern for TAEST16001.

### 3.6 Assessment of anti-tumor efficacy of TAEST16001 in xenograft models

To determine *in vivo* anti-tumor efficacy of TAEST16001, we employed a series of human tumor xenograft models. Figure 5A illustrates the overall process of the experiments. In the first model, NOD/SCID/IL2gR^-/-^ (NSG) mice were subcutaneously engrafted with human NSCL cell line NCI-H1299 overexpressing HLA-A2 (NCI-H1299-A2) and treated with different doses of TAEST16001. Tumor growth was significantly inhibited by treatment with all doses of TAEST16001 (figure 5B). Tumor growth was nearly completely inhibited at higher dosages (1× 10^7^ and 2 × 10^7^ cells per mouse, figure 5B). In a control experiment, TAEST16001 failed to inhibit growth of HLA-A2 negative NCI-H1299 wild-type tumors (online supplemental figure 5), suggesting that tumor inhibition was antigen-specific. Similar tumor regression by TAEST16001 treatment was also observed in fibrosarcoma (online supplemental figure 6A) and melanoma (online supplemental figure 6B) models. Moreover, immunohistochemical studies revealed the extensive presence of CD8^+^ cells in the tumor microenvironment in mice in the TAEST16001 group but not in the control TCR-T group (figure 5C).

**Figure 5.**
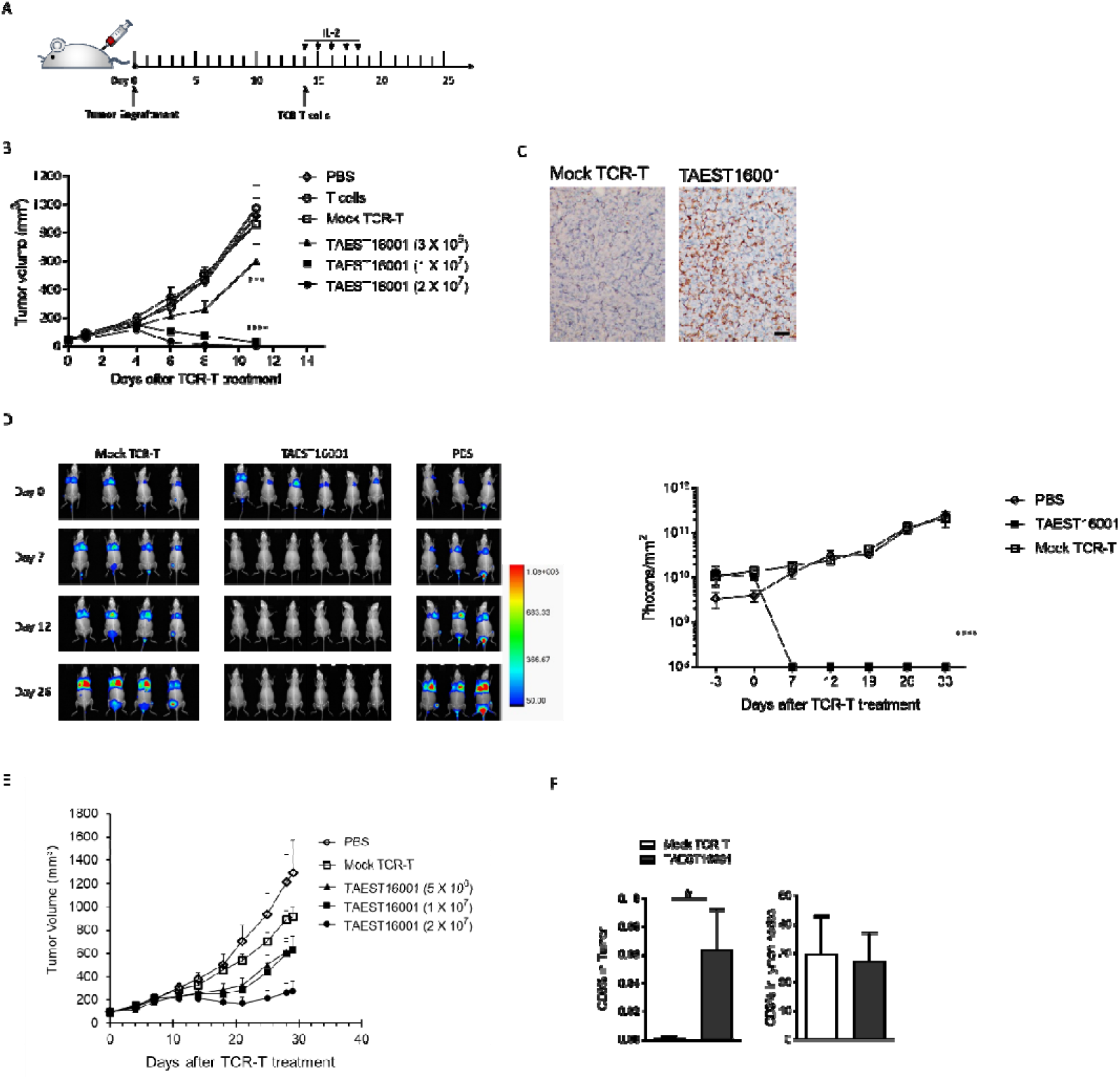
TAEST16001 shows high anti-tumor efficacy *in vivo.* (A) Schematic representation of TAEST16001 treatment of human tumor xenograft models. Tumor cells were engrafted in NSG mice. After establishing the tumor, mice were treated with TCR-T cells, followed by five consecutive injections of IL-2. The tumors were monitored regularly after treatment. (B) TAEST16001 inhibited tumor growth in a xenograft model of lung cancer. NSG mice engrafted with NCI-H1299-A2 NSCLC cells were treated with indicated doses of TAEST16001. Vehicle (PBS), T cells without TCR transduction (T cells, 2 × 10^7^), and T cells transduced with A6 TCR (Mock TCR-T, 1 × 10^7^) were used as controls. N = 5 mice per group. At doses of 1 × 10^7^ and 2 × 10^7^, the differences between TAEST16001 and the Mock TCR-T were highly significant from day 6 to day 11 (****, P < 0.0001, two-way ANOVA). At doses of 3 × 10^6^, the difference between TAEST16001 and Mock TCR-T were highly significant from day 8 to day 11 (***, P < 0.001, two-way ANOVA). (C) TAEST16001 cells but not control TCR-T cells infiltrated in the tumor microenvironment. NSG mice engrafted with NCI-H1299-A2 NSCLC cells were treated with 6 × 10^6^ TAEST16001 or control TCR-T cells. Forty-eight hours post treatment, tumor sections were collected and stained for human CD8 and analyzed using immunohistochemical staining. Scale bar = 50 μm. (D) TAEST16001 inhibited metastasis. NSG mice implanted with A549^Luciferase/A0201/NY-ESO-1^ cells were treated with 1 × 10^7^ TAEST16001 or mock TCR-T. Imaging data from day 0 to day 26 (left) and quantitative analysis of photon counts (right) are shown here. The difference between TAEST16001 and the Mock TCR-T were highly significant (****, P < 0.0001, two-way ANOVA) from day 7 to day 33. (E) TAEST16001 inhibited tumor growth in a NSCLC PDX model. NSG mice engrafted with PDX tumors were treated with indicated doses of TAEST16001 or control TCR-T cells. N = 8 mice per group. (F) TAEST16001 infiltrated the PDX tumor. NSG mice engrafted with PDX tumors were treated with 5 × 10^6^ TAEST16001 or mock TCR-T cells. At the end of the experiments, tumors and lymph nodes were collected, and the percentage of CD3^+^ cells in total cells was calculated, *, P < 0.05, Student’s t-test.

To determine whether TAEST16001 can inhibit metastasis, lung metastasis model was established by tail vein injection of A549 cells (overexpressing luciferase, HLA-A2, and NY-ESO-1), and the mice were treated with 1 × 10^7^ TAEST16001 cells. No tumor growth was observed in any of the six TAEST16001-treated mice, whereas all mice treated with PBS or control TCR-T developed lung metastasis (figure 5D).

Furthermore, we analyzed the anti-tumor efficacy of TAEST16001 in a patient-derived xenograft (PDX) model. PDX models preserve the heterogeneity and microenvironment of human tumors and thus are more clinically relevant than the tumor cell line models studied above. NSG mice engrafted with NSCLC PDX tumors (HLA-A2^+^ and NY-ESO-1^+^) were treated with different doses of TAEST16001. Significant inhibition of PDX tumor growth was observed, especially at high doses of 2 × 10^7^ cells per mouse (figure 5E). Flow cytometry analysis revealed that significantly more T cells infiltrated the tumor in the TAEST16001 treated group than in the control TCR-T group. In contrast, no significant difference in the lymph nodes was observed in the mice of the two groups (figure 5F). Our data suggested that TAEST16001 could effectively infiltrate the tumor microenvironment and inhibit PDX tumor growth.

## 4 DISCUSSION

The recent approval of KIMMTRAK (tebentafusp), a novel TCR/anti-CD3 bispecific fusion protein targeting gp100, for the treatment of metastatic uveal melanoma^35^ is a historic breakthrough. It paved the way for the development of effective TCR-based immunotherapies for solid tumors. However, the advancement of TCR-T therapy is hindered by the lack of a development platform to bring safe and effective TCR-T products to clinical trials. Traditional drug discovery processes are no longer suitable for TCR-based therapies. In this study, we detailed a development platform combining T cell cloning, TCR engineering, efficacy testing and safety screening techniques. This robust platform allowed the successful development of TAEST16001, which is under clinical investigation in a phase I trial (NCT03159585).

The first step towards developing TCR-engineered T cell therapies is to obtain TAA-specific TCRs. To date, most therapeutic TCRs targeting TAAs used in ACT clinical trials have been derived from tumor-infiltrating T cells (TILs) of resected tumors^21 36^ or peripheral blood of vaccinated patients.^17^ Although patient-derived TCRs are effective, they are limited by the availability of proper tumor patients. A more convenient approach is to isolate TCRs from the peripheral blood of immunized transgenic mice.^36 37^ However, safety concerns of the murine TCRs, including potential immune response to the xenogeneic proteins in patients^38^ and cross-reactivity due to the lack of thymic selection in humans, should not be overlooked. Here, we decided to acquire TCRs specific for TAAs directly from the PBMCs of HLA-matched healthy donors. Contrary to popular belief that T cells reactive with self-proteins are eliminated by the clonal deletion in healthy humans, it has been demonstrated that thymic selection does not eliminate as much as prune self-specific T cells.^39^ Therefore, PBMCs of healthy donors are convenient and reliable sources of TAA-specific TCRs.

TCRs engage with pMHC ligands through three CDRs: germ line-encoded CDR1 and CDR2 and somatically rearranged CDR3. Structural studies have revealed that, in general, CDR1 and CDR2 make primary contact with the MHC surface, whereas CDR3 interacts with the peptide epitope.^40^ Mutations can generate high-affinity TCRs in all three CDRs, ^27 41 42^ and the combination of mutations of different CDR3 can generate TCRs with picomolar affinity.^27^ Although CDR1 and CDR2 are situated close to the MHC, mutations in CDR1 and CDR2 can generate high-affinity TCRs without sacrificing specificity. The gain in affinity has been attributed mainly to the improved shape complementarity of the CDRs, rather than direct contact with pMHC.^42^ However, because of the binding geometry between TCR and pMHC, mutations in CDR1 and CDR2 increase the likelihood of TCR engaging with the helical regions of MHC and thus reduce peptide specificity; mutations on CDR3, on the other hand, may promote interactions with the peptide and consequently increase peptide specificity. Because TCRs with optimal functional avidity only require moderate affinities, and in most cases, mutations on one CDR will suffice. Therefore, we focused only on CDR3, which is the safest from a structural point of view.

Generally, TCRs with affinities in the range of 1-10 μM show optimal functional avidities.^9 14 17^ SL-A10B0 has an affinity of ~1.5 μM, which is within this range. However, other SL3 or SL2 mutants with similar affinities showed non-optimal avidity or even cross reactivity, indicating that factors other than binding affinity can also contribute to TCR function. First, accumulating evidence suggests involvement of structural mechanisms. The type of bond between TCR and pMHC interface determines the functional outcomes:^43 44^ catch bonds (i.e., dissociation lifetime extends under force) favor T cell activation, while slip bonds (i.e., dissociation lifetime decreases with increasing force) cause non-responsiveness. Docking geometry between TCR and pMHC also plays a critical role in determining TCR functional outcomes. Deviation from the stereotypical docking geometry tends to limit TCR signaling.^45 46^ Second, SPR assay determines the binding between proteins in three-dimensional (3D) solution (3D binding), while in reality both TCR and pMHC are anchored on twodimensional (2D) cell membranes (2D binding). 2D binding kinetics are dramatically different from 3D binding kinetics, and also better correlate with TCR functions.^47 48^ Thus, the complexity of TCR/pMHC interaction makes it difficult to predict TCR function. Screening a large panel of TCR mutants is a preferred strategy for selection of a lead candidate with optimal potency and specificity.

In order to cope with the vast amount of peptide epitopes using a limited TCR repertoire in the immune system, each TCR must cross-react with multiple peptides.^49^ Cross-reactivity is not a matter of concern to TCR-T therapies unless TCRs unexpectedly recognize antigens expressing in normal tissues, leading to toxicity in clinical trials.^15 16^ TCR-pMHC binding modes determine TCR cross-reactivity: TCRs making more contacts with peptide side chains exhibit a lesser degree of cross-reactivity.^50^ In this study, we investigated the binding modes of SL3-A10B0 and 1G4-α95:LY using alanine-scanning mutagenesis and found that 1G4-α95:LY binding to NY-ESO-1 _157-165_ was dominated by one residue (position 5), whereas four residues (positions 3, 5, 6 and 7) were critical for SL3-A10B0 binding. Therefore, we anticipate that SL3-A10B0 will have a better safety profile in clinical trials.

In conclusion, TAEST16001 has demonstrated superior efficacy against NY-ESO-1/HLA-A2 tumors and an excellent safety profile in our extensive *in vitro* and *in vivo* experiments. The development strategy presented here can be applied to any affinity-enhanced TCR-T cells and greatly expands the opportunities for TCR-T therapies.

## Acknowledgements

We would like to thank Yi Li, Lun Zeng and Yonghui Wang for administrative supports. We would like to thank Wenzhuo Zhao, Danfeng Shi, Ning Wang, Yanping Pan, Weilin Wang, Congming Fang, Mengyong Yan and Yaolong Chen for technical supports. We would like to thank Editage (www.editage.cn) for English language editing.

