## Supplemental Table for "Development of an affinity-enhanced clinical candidate TCR targeting NY-ESO-1 with optimal potency and high specificity"

Development of affinity-enhanced TCR-modified T cells targeting NY-ESO-1 for adoptive cell therapy

### Supplemental

#### Supplemental Table 1 The expression levels of NY-ESO-1 (encoded by *CTAG1*) and NY-ESO-2 (encoded by *CTAG2*) in tumor cell lines were assessed using the NanoString nCounter system.

| Cell Line | HLA-A | Number of Counts | |
| --- | --- | --- | --- |
|  |  | *CTAG1* | *CTAG2* |
| A375 | A*01:01/A*02:01 | 3525 | 20 |
| IM9 | A*02:01/A*02:05 | 189 | 498 |
| U266B1 | A*02:01/A*03:01 | 6940 | 4024 |
| K562 | A*11:01/A*31:01 | 4 | 516 |
| MDA-MB-231 | A*02:01/A*02:17 | 1 | 9 |
| MEL526 | A*02:01/A*03:01 | 4 | 5 |
| NCI-H1299 | A*24:02/A*32:01 | 5357 | 13 |
| NCI-H522 | A*02:01/A*02:01 | 1 | 4602 |
| NCI-H1650 | A*02:01/A*02:01 | 2 | 3 |

#### Supplemental Table 2 Peptides derived from normal proteins

| **peptide** | **Protein** | **Gene** | **Position** | **Affinity** |
| --- | --- | --- | --- | --- |
| FLLEKGYEV | GDP-mannose 4,6 dehydratase | *GMDS* | 42-50 | no binding |
| ALLDRIVSV | Nuclear pore complex protein Nup205 | *NU205* | 1498 - 1506 | no binding |
| FLLDKKIGV | T-complex protein 1 subunit beta | *TCPB* | 218 - 226 | no binding |
| TLWVDPYEV | Protein BTG1 | *BTG1* | 103 - 111 | no binding |
| KLFGMIITI | Protein transport protein Sec61 subunit alpha isoform 1 | *S61A1* | 117 - 125 | no binding |
| FLPPLPTSV | Bifunctional coenzyme A synthase | *COASY* | 96 - 104 | no binding |
| FLDPRPLTV | Cytochrome P450 1B1 | *CP1B1* | 190 - 198 | no binding |
| ILWETVPSM | Fibronectin type III domain-containing protein 3B | *FND3B* | 1072 - 1080 | no binding |
| VLLGKVYVV | Kelch-like protein 24 | *KLH24* | 404 - 412 | no binding |
| SLEDILHQV | Fibrinogen gamma chain | *FIBG* | 68 - 76 | no binding |
| FVFPGELLL | Neutral amino acid transporter B | *AAAT* | 89 - 97 | no binding |
| SLLGGDVVSV | TSC22 domain family protein 3 | *T22D3* | 27 - 36 | no binding |
| ALLKYIETL | ATP-dependent RNA helicase A | *DHX9* | 639 - 647 | no binding |
| ILTDITKGV | Elongation factor 2 | *EF2* | 661 - 669 | no binding |
| TLWGIQKEL | L-lactate dehydrogenase A chain | *LDHA* | 322 - 330 | no binding |
| RLDELGGVYL | Dolichyl-diphosphooligosaccharide--protein glycosyltransferase subunit 2 | *RPN2* | 190 - 199 | no binding |
| FVNDIFERI | Histone H2B type 1-B | *H2B1B* | 66 - 74 | no binding |
| LLGPPPVGV | Cip1-interacting zinc finger protein | *CIZ1* | 159 - 167 | no binding |
| SLLMWITQC | Cancer/testis antigen 1 | *CTG1B* | 157 - 165 | 1.77 M |

#### Supplemental Table 3 Binding properties of SL3-A10B0 and 1G4-α95:LY to HLA-A2 in complex with alanine substitutions of the NY-ESO-1_157-165_ peptide. The binding was determined by SPR.

| **Mutant** | **SL3-A10B0** | | | **1G4-α95:LY** | | |
| --- | --- | --- | --- | --- | --- | --- |
|  | *k_a_* (1/Ms) | *k_d_* (1/s) | *K*_D_ (M) | *k_a_* (1/Ms) | *k_d_* (1/s) | *K*_D_ (M) |
| **WT** | 1.131E+05 | 1.640E-01 | 1.449E-06 | 3.013E+04 | 3.916E-02 | 1.300E-06 |
| **S1A** | 1.209E+05 | 1.758E-01 | 1.454E-06 | 2.855E+04 | 4.378E-02 | 1.533E-06 |
| **L3A** | 3.957E+03 | 1.657E+00 | 4.188E-04 | 3.448E+04 | 8.150E-02 | 2.364E-06 |
| **M4A** | 7.901E+03 | 3.547E-01 | 4.489E-05 | 6.358E+03 | 1.008E+00 | 1.585E-04 |
| **W5A** | No binding | No binding | No binding | No binding | No binding | No binding |
| **I6A** | 2.288E+03 | 1.269E+00 | 5.547E-04 | 5.941E+03 | 3.646E-01 | 6.137E-05 |
| **T7A** | 2.739E+03 | 1.677E+00 | 6.122E-04 | 1.042E+04 | 1.830E-01 | 1.756E-05 |
| **Q8A** | 2.326E+05 | 1.093E-01 | 4.698E-07 | 1.083E+04 | 1.280E+00 | 1.182E-04 |
| **C9A** | 1.367E+05 | 1.056E-01 | 7.726E-07 | 3.050E+04 | 4.531E-02 | 1.486E-06 |

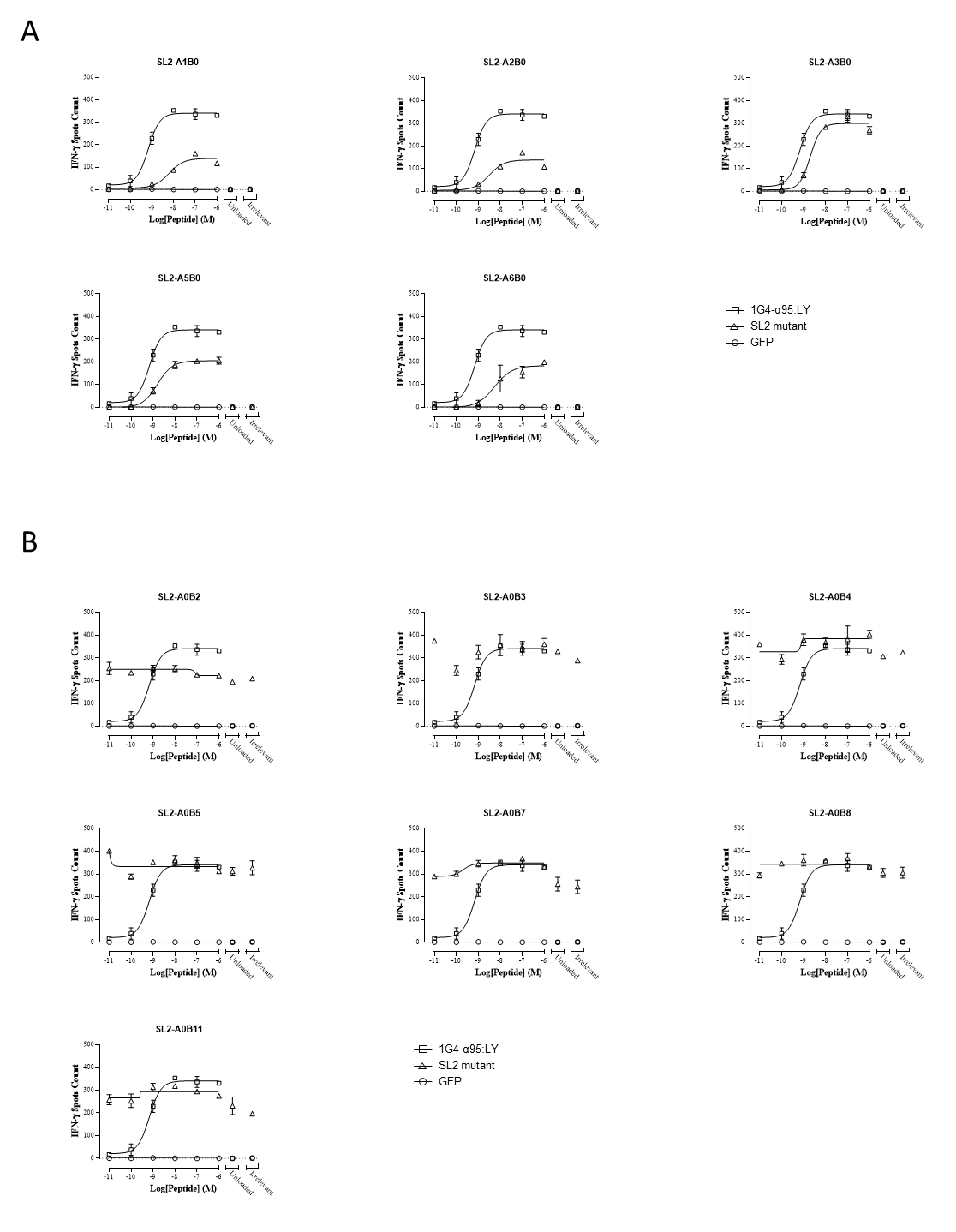

#### Supplemental Figure 1 Functional activities of affinity-enhanced SL2 mutants. IFN-γ release by CD8^+^ T cells transduced with affinity-enhanced SL2 α-chain (A) or β-chain (B) mutants in the presence of T2 cells pulsed with various concentrations of NY-ESO-1_157-165_, an irrelevant peptide (10^-6^ M gp100_280-288_, YLEPGPVTA), or no peptide (Unloaded). T cells expressing GFP served as the negative control and 1G4-α95:LY as a positive control.

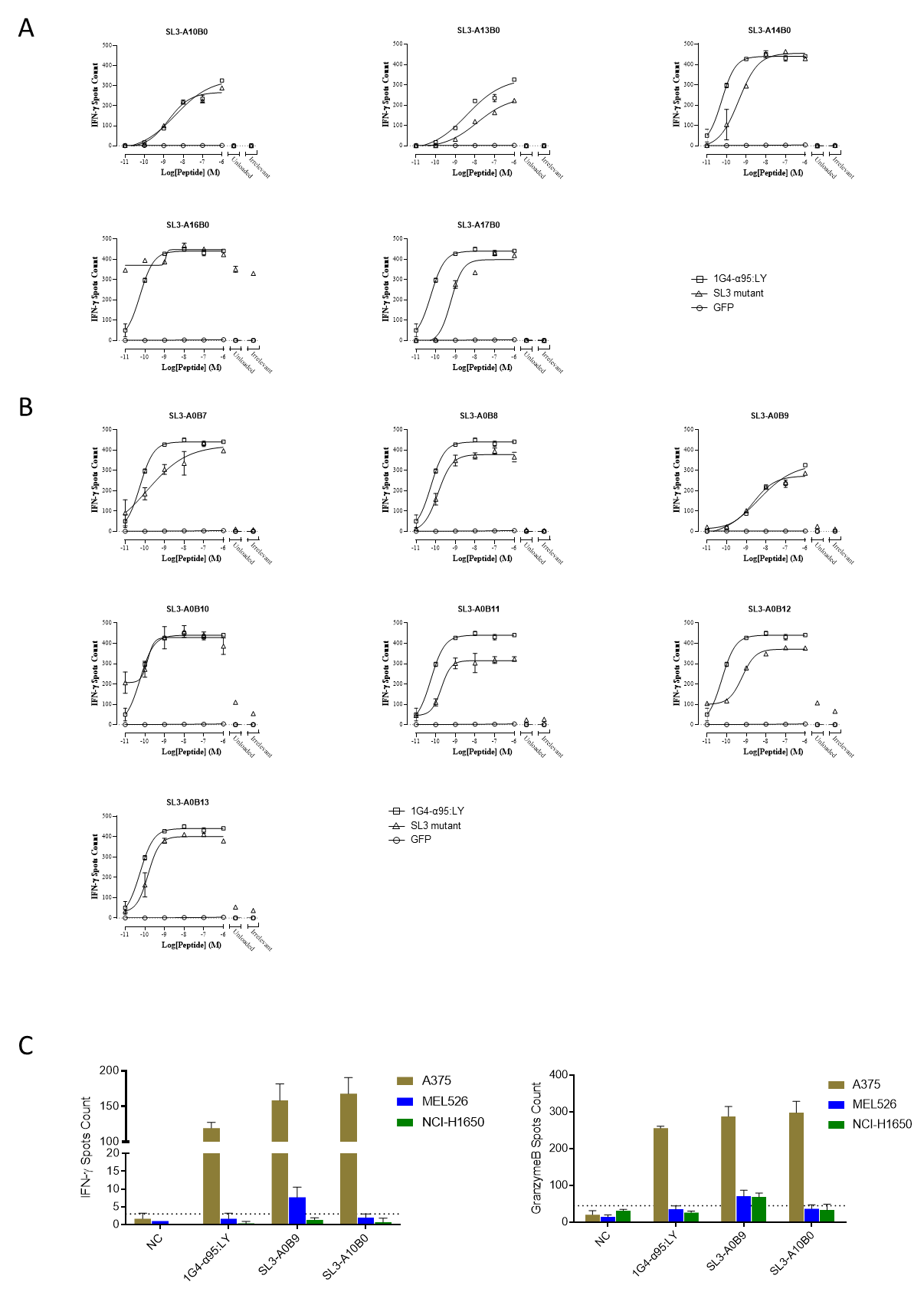

#### Supplemental Figure 2 Functional activities of affinity-enhanced SL3 mutants. IFN-γ release by CD8^+^ T cells transduced with affinity-enhanced SL3 α-chain (A) or β-chain (B) mutants in the presence of T2 cells pulsed with various concentrations of NY-ESO-1_157-165_, an irrelevant peptide (10^-6^ M gp100_280-288_, YLEPGPVTA), or no peptide (Unloaded). T cells expressing GFP served as the negative control and 1G4-α95:LY as a positive control. （C）Non-specific activation of SL3-A0B9. IFN-γ (left panel) or Granzyme B (right panel) release of CD8^+^ T cells expressing SL3-A0B9, SL3-A10B0 or 1G4-α95:LY after co-culturing with tumor cell lines: A375 (HLA-A2^+^, NY-ESO-1^+^/NY-ESO-2^-^), MEL526 (HLA-A2^+^, NY-ESO-1^-^/NY-ESO-2^-^) and NCI-H1650 (HLA-A2^+^, NY-ESO-1^-^/NY-ESO-2^-^). Non-transduced T cells (NT) served as a negative control to assess background activation levels (dotted lines).

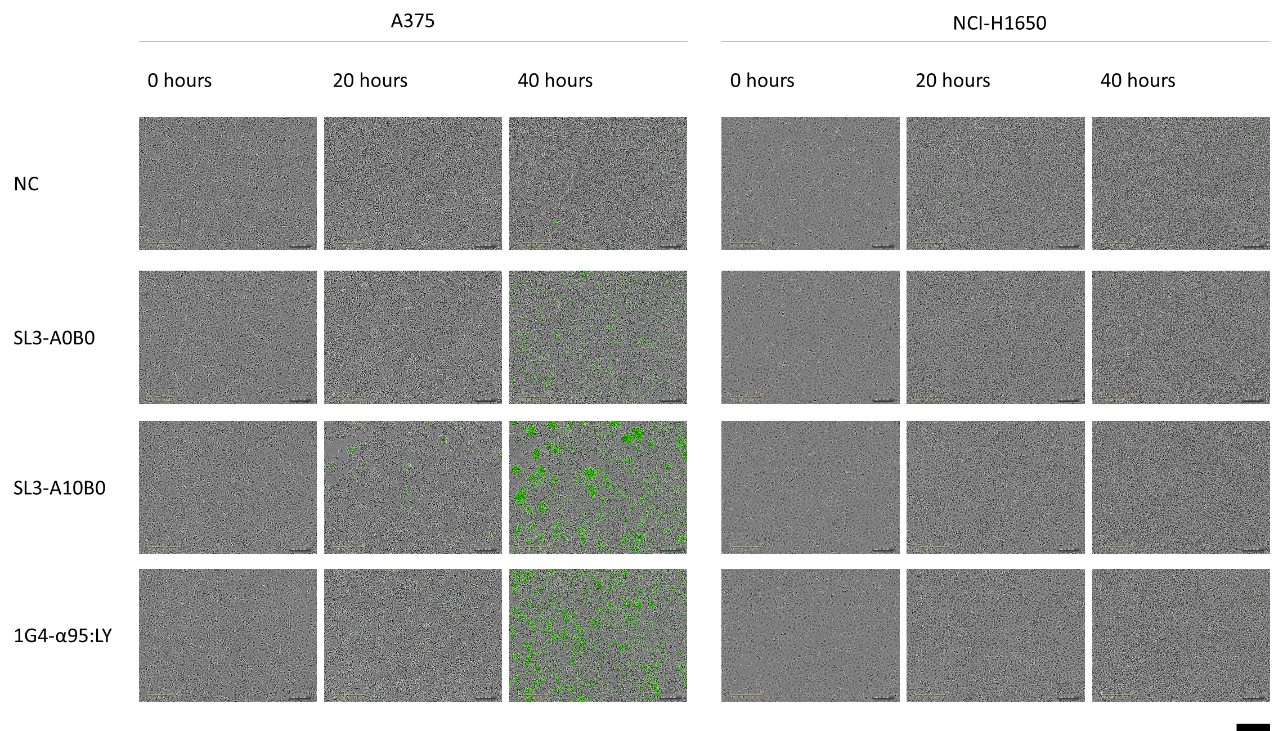

#### Supplemental Figure 3 Representative images of the live-imaging killing assay (figure 2C). Overlay of the bright-field and green fluorescence images at 0, 20 and 40 hours. Apoptotic tumor cells were labeled green. Scale bar = 300 μm.

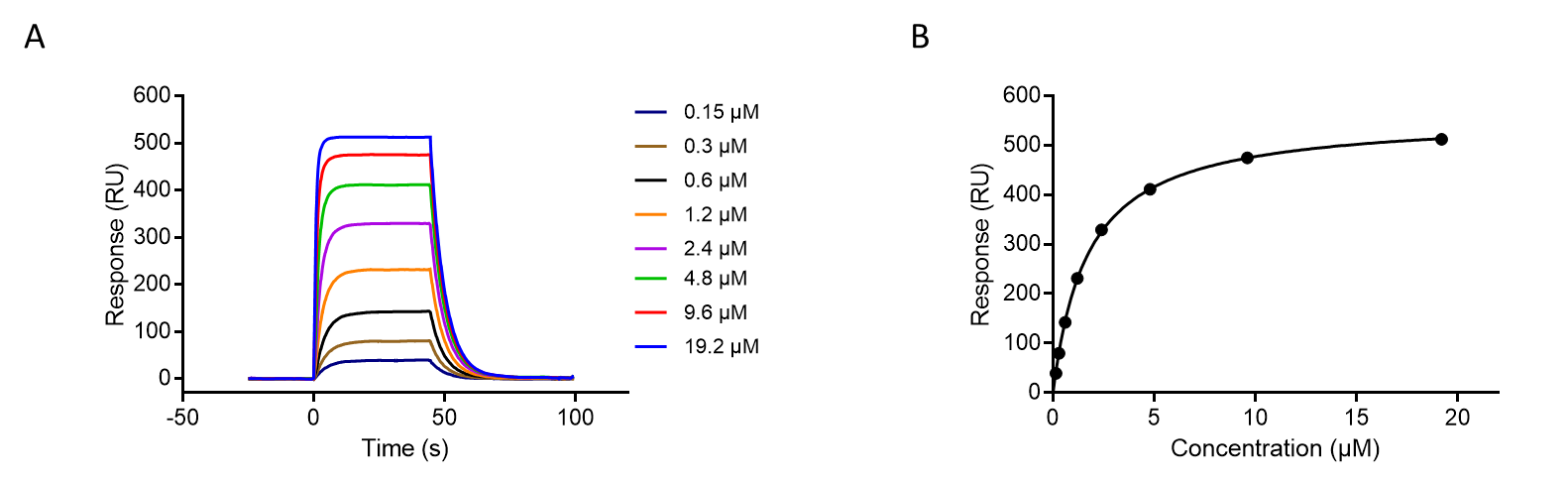

#### Supplemental Figure 4 SPR analysis of SL3-A10B0. (A) Sensograms of kinetic measurement of SL3-A10B0 TCR to its cognate pMHC ligand NY-ESO-1_157-165_/HLA-A2. Soluble TCRs at the indicated concentrations were sequentially injected to biotinylated pMHC immobilized on a streptavidin-coated sensor chip surface. (B) The responses at equilibrium binding were fitted with the 1:1 Langmuir model to determine the equilibrium dissociation constant (*K*_D_).

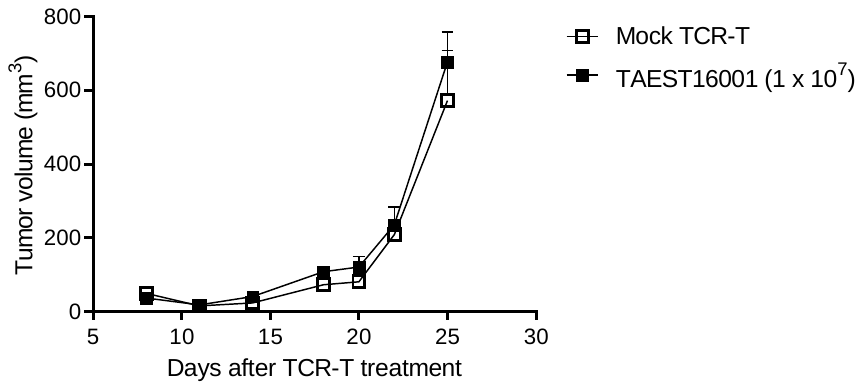

#### Supplemental Figure 5 TAEST16001 did not inhibit tumor growth in an HLA-A2^-^ xenograft model. NSG mice engrafted with NCI-H1299 cells (HLA-A2^-^, NY-ESO-1^+^) were treated with the indicated dose of TAEST16001. T cells transduced with A6 TCR (Mock TCR-T) were used as a negative control. N = 5 mice per group.

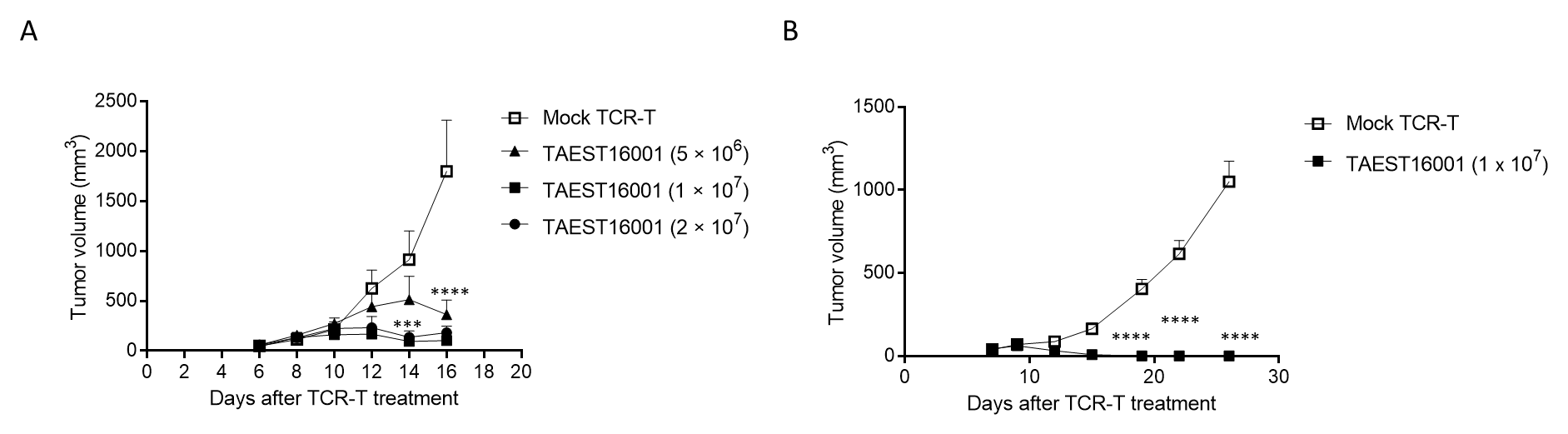

#### Supplemental Figure 6 TAEST16001 inhibited tumor growth in fibrosarcoma (A) and melanoma (B) xenograft models. (A) NSG mice engrafted with HT1080 fibrosarcoma cells overexpressing HLA-A2 (HT1080-A2) were treated with indicated doses of TAEST16001. T cells expressing A6 TCR (Mock TCR-T, 1 × 10^7^) served as a negative control. N = 5 mice per group. On day 14, the differences between TAEST16001 and Mock TCR-T were highly significant at doses of 1 × 10^7^ and 2 × 10^7^ (***, P < 0.001, two-way ANOVA). On day 16, the differences between TAEST16001 and Mock TCR-T were highly significant at all doses (****, P < 0.0001, two-way ANOVA). (B) NSG mice engrafted with A375 melanoma cells were treated with TAEST16001 (1 × 10^7^ per mouse). T cells expressing A6 TCR (Mock TCR-T, 1 × 10^7^) served as a negative control. N = 5 mice per group. The differences between TAEST16001 and Mock TCR-T were highly significant from day 20 to 26 (****, P < 0.0001, two-way ANOVA).
